# OSDR2.0 infers microenvironment-driven cell-state transitions and population dynamics from a single spatial biopsy

**DOI:** 10.1101/2025.09.02.673328

**Authors:** Itay Ben Shalom, Jonathan Somer, Shoval Miyara, Avi Mayo, Shie Mannor, Uri Alon

## Abstract

Cell populations in human tissues change over time by cell division, death and transitions between functional states. In the tumor microenvironment (TME), such dynamics are central to immune evasion, stromal remodeling and therapeutic response. However, it is difficult to measure such dynamics in vivo because usually only a single biopsy is available providing a static snapshot. To obtain cell population dynamics from a snapshot we previously developed One Shot Dynamic Reconstruction (OSDR1.0), an algorithm that reconstructs cell population dynamics over weeks to months from a spatial biopsy using cell-type information and a proliferation marker. OSDR1.0 however does not include transitions between cell states. Here we present OSDR2.0, an extension that incorporates transitions between cell states inferred from the local cellular neighborhood. The algorithm OSDR2.0 models the probability of a cell being in a given state (e.g., PD1^+^ vs. PD1^−^ T cell, or cancer-associated vs. resting fibroblast) as a function of its surrounding cell types. These state-transition rules are then integrated into simulations of population dynamics. After the cell population is advanced by a timestep, the cell states are adjusted according to the new neighborhoods, using the fact that cell state transitions, which take hours, are typically much more rapid than cell population changes on the scale of weeks, and can thus be considered at quasi-steady-state. Applying OSDR2.0 to spatial proteomics data from triple-negative breast cancer (TNBC), we find that cell state is strongly associated with local microenvironment composition. Incorporating state transitions significantly improves the ability to predict treatment response - OSDR2.0 accurately separates responders from nonresponders to chemotherapy and immunotherapy based on early post-treatment biopsies, outperforming OSDR1.0 that does not include cell state transitions. This work highlights the importance of cell state plasticity in shaping tumor response and opens a way to infer both population dynamics and cell state transitions from static clinical samples.

## Introduction

Understanding how cells interact and change over time in human tissues is central to uncovering the principles of development, immunity, and disease progression (Meizlish *et al*., 2021; Inayatullah, Dwivedi and Tiwari, 2025; Zhang *et al*., 2025). In cancer, dynamic interactions between tumor cells and their surrounding microenvironment, including immune and stromal cells, play a key role in shaping tumor evolution and treatment response (Gascard and Tlsty, 2016; Binnewies *et al*., 2018; Sahai *et al*., 2020). These processes involve not only changes in the abundance of different cell types, but also rapid transitions between functional states within a given cell type, such as from resting to activated or exhausted phenotypes in T cells (Giles *et al*., 2022; Rudloff *et al*., 2023), or from resting fibroblasts to cancer-associated fibroblasts (CAFs) (Lavie *et al*., 2022; Mellone *et al*., 2022). Capturing these complex dynamics remains a major challenge, especially in human samples where time-resolved measurements are generally unavailable.

We recently introduced One-Shot Dynamics Reconstruction (OSDR, which we call here OSDR1.0) (Somer, Mannor and Alon, 2025), an algorithm that infers cell population dynamics from a single spatial biopsy by using cell-type annotations and proliferation markers to estimate division and removal rates as a function of local neighborhood composition. OSDR1.0 uncovered dynamic tissue circuits, such as bistable macrophage-fibroblast feedback and excitable T/B-cell interactions, and predicted treatment responses in breast cancer from early-treatment snapshots alone.

However, OSDR1.0 treats each cell type as a homogeneous population and does not account for transitions between cell states. This neglects potentially important mechanisms in which cell states, such as Programmed Cell Death Protein 1 (PD1) expression in T cells, can shift rapidly and are crucial determinants of immune function and therapeutic efficacy (Jerby-Arnon *et al*., 2018; Schürch *et al*., 2020; Giles *et al*., 2023).

Here we present *OSDR2.0*, an extension of the OSDR framework that incorporates cell state dynamics into spatial modeling. OSDR2.0 learning the probability of a cell to be in a particular state (e.g., PD ^+^ vs. PD ^−^ T cells or CAF vs. resting fibroblast) given the composition of its cellular neighborhood. OSDR 2.0 leverages the fact that cell state transitions typically occur on a faster timescale than cell proliferation - cell state changes (such as T cell exhaustion and fibroblast activation) typically take on the order of hours to days (Manella *et al*., 2020; Kirschenbaum *et al*., 2024), whereas significant changes in cell population take weeks even in rapidly growing cancers (Kay *et al*., 2019; Narayanan *et al*., 2025). During dynamic simulations, the model first updates cell division and removal, and then adjusts the cell state according to the new neighborhoods. This enables OSDR 2.0 to infer not only the trajectories of cell-type population abundances over time, but also the evolving distributions of functionally distinct cell states within those populations. Here we demonstrate this on two cell states, and note that the same approach can be generalized to multiple cell states in the same cell type.

Applying OSDR2.0 to spatial proteomics data from triple-negative breast cancer (TNBC) (Wang *et al*., 2023), we find that cell states are strongly correlated with local neighborhood composition. Including these state transitions markedly improves the model’s ability to predict patients’ response to therapy. In particular, OSDR2.0 separates responders from nonresponders to chemotherapy and immunotherapy with higher accuracy than models that consider cell types abundance alone. By integrating rapid intra-cell-type plasticity with long-timescale population dynamics, OSDR2.0 offers a nuanced and predictive view of tissue behavior from a single biopsy snapshot, with potential applications across diverse tissues and pathologies.

## Results

### OSDR2.0 estimates dynamics of cell states and populations based on a spatial snapshot

We build on OSDR1.0 that estimates cell population dynamics over weeks from a tissue biopsy (Somer, Mannor and Alon, 2025). The input data is a spatial proteomics snapshot that includes cell type identity and a cell division marker (Fig 1a). The OSDR1.0 algorithm learns the division and removal rates of a cell type, as a function of its neighborhood composition. Using the inferred division and removal rates, OSRD1.0 simulates cell division and removal of cells, updates cellular neighborhoods accordingly, and repeats.

**Figure 1.**
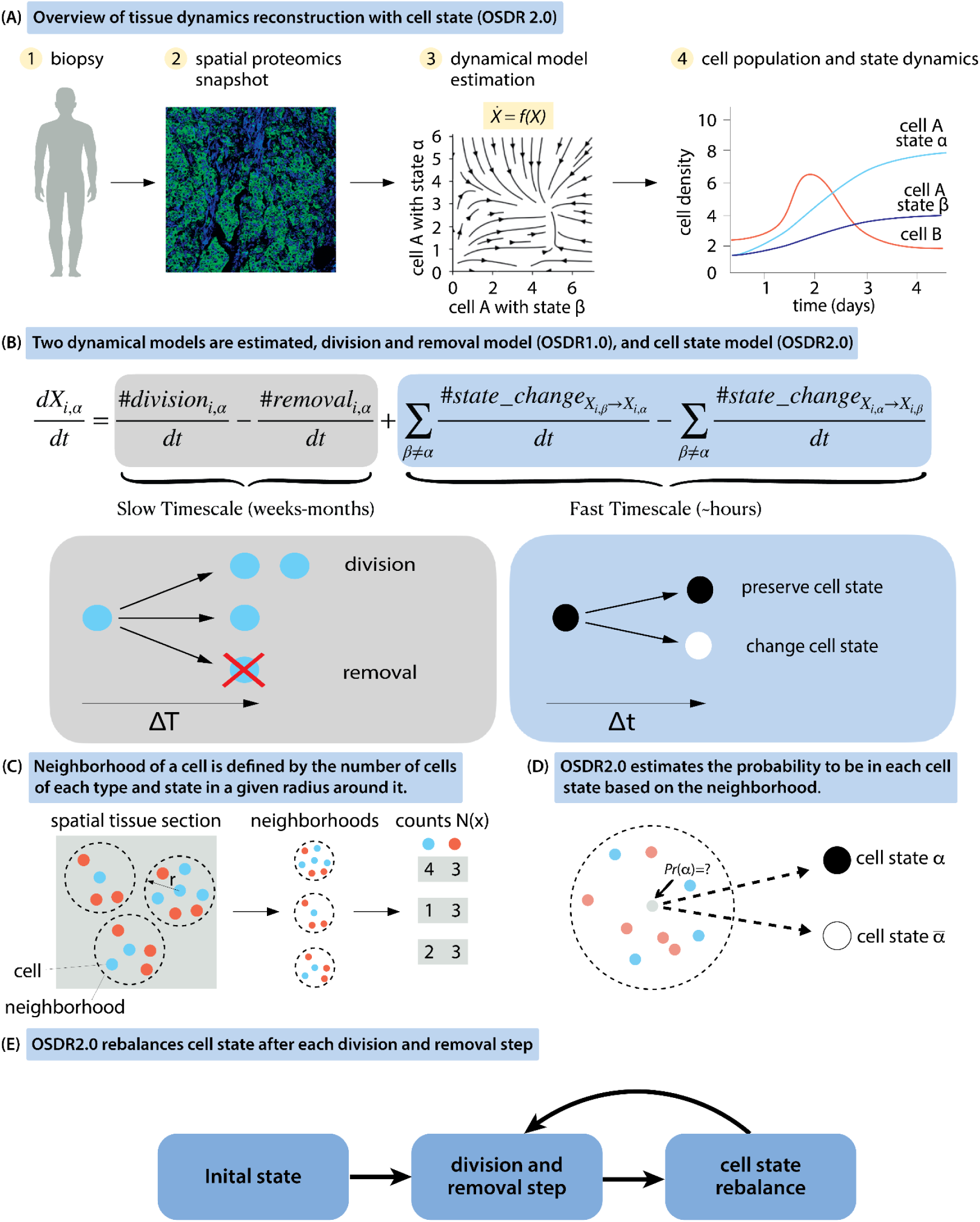
Overview of the OSDR2.0 approach to estimate cell state dynamics and population dynamics from a tissue biopsy. (A) OSDR2.0 uses as input a spatial omics dataset of cell positions, types and states, along with a division marker (e.g. Ki67). It learns division and state probabilities as a function of neighborhood composition to allow for phase portrait analysis and simulations of cell populations and states over weeks. (B) OSDR2.0 adds transitions between cell states to OSDR1.0. A cell type 𝑋_*i*_ that has several states, 𝑋_*i*,1_, 𝑋_*i*,2_, … denoted 𝑋_*i*,α_. The rate of change of the cell state populations is given by division and removal and the balance of transitions between states. The model uses the fact that cell-state transitions are typically much faster than the division and removal processes. (C) A cell’s neighborhood is defined by the counts of cells of each type and state in a radius r around it, in this study r=80um. (D) OSDR2.0 estimates the probability of each cell type to be in each of its states as a function of the neighborhood composition. (E) OSDR2.0 simulates cell dynamics by starting with the initial condition of cells in space, simulating divisions and removal according to each cell’s neighborhood, and adjusting cell sets according to the new neighborhoods. This is repeated to form temporal trajectories. Figure adapted from Somer (Somer, Mannor and Alon, 2025).

Here we add to OSDR1.0 a method to compute cell states (Fig 1b). Cell states are functionally distinct, and are classically defined by the expression of marker genes and proteins, for example with T-cells one can define PD ^+^ and PD ^−^ T cells where the first cell state is known to exhibit impaired cytotoxic activity compared to the latter (Ahmadzadeh *et al*., 2009; Balança *et al*., 2021).

The new algorithm, OSDR2.0, takes the neighborhood of each cell (Fig 1c) and its cell-state, and learns the probability that a cell type is in state α or state ᾱ as a function of the neighborhood composition of cell types (without cell states). OSDR2.0 learns the probability function by fitting a logistic regression model (Fig 1d).

OSDR2.0 then uses the fact that cell state transitions are typically much faster than cell population changes due to proliferation and removal.Thus, OSDR2.0 starts with an initial cell spatial configuration, evaluates division and removal to update the cell types. It then uses separation of timescales, by adjusting cell states according to the new neighborhood compositions (Fig 1e). This enables OSDR2.0 to follow both cell population dynamics and cell state dynamics.The OSDR2.0 regression coefficients indicate which cell types in the neighborhood affect cell division and cell states, inhibiting or enhancing each state and at which relative strength.

### Cell state is strongly associated with neighborhood cell-type composition

The OSDR2.0 method relies on the assumption that neighborhood composition is a strong correlate of cell state (Keren *et al*., 2018; Hickey *et al*., 2022; Ding *et al*., 2025; Somer, Mannor and Alon, 2025). To test this, we analyzed two breast cancer imaging mass cytometry (IMC) datasets (Danenberg *et al*., 2022; Wang *et al*., 2023) which provide spatially resolved single-cell type, division marker and state markers. The datasets include major immune and stromal cell types, across multiple patient samples. We focused on two biologically relevant state transitions in the tumor microenvironment: (i) fibroblasts transitioning between a resting state and CAFs (Fig. 2a,c), and (ii) T cells transitioning between PD1⁻ and PD1⁺ states (Fig. 2b,d). These transitions are central to tumor progression: CAF activation is associated with extracellular matrix remodeling (Barbazan *et al*., 2023), immune suppression (Tsoumakidou, 2023), and enhanced tumor invasiveness (Caligiuri and Tuveson, 2023), and are considered a therapeutic target (Cao *et al*., 2025). PD1 expression on T cells reflects an exhausted phenotype that impairs anti-tumor immunity and serves as a target for immune checkpoint blockade therapies (Ahmadzadeh *et al*., 2009; Balança *et al*., 2021; Giles *et al*., 2023).

**Figure 2.**
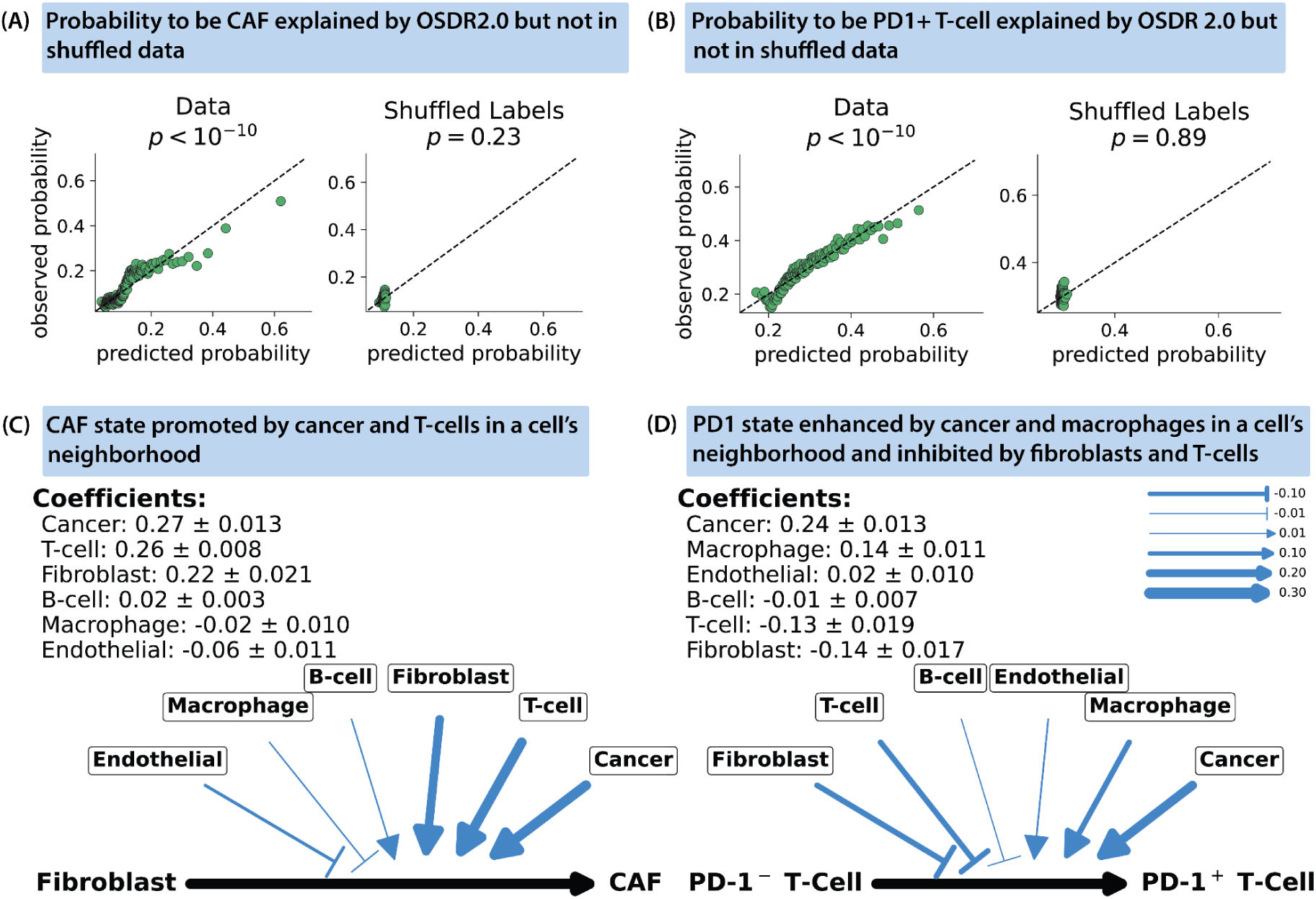
Cell state is strongly correlated to neighborhood composition. (A) Probability to be CAF versus fibroblast in data and OSDR2.0 prediction are highly correlated. The correlation between shuffled data and model prediction is poor. (B) Probability to be PD1^+^ T cell versus PD1^−^ T cell in data and OSDR2.0 predictions are highly correlated. The correlation between shuffled data (where PD1^+^/PD1^−^ labels were randomly shuffled) and model prediction is poor. In A-B, each dot is a bin of 1000 cells and the p-value is computed for the model using likelihood ratio test using statmodels (Seabold and Perktold, 2010). (C) Regression coefficients in OSDR2.0 model for the probability to be CAF versus fibroblast shows strong effects of cancer and fibroblast cells (all 95% confidence intervals do not include 0). (D) Regression coefficients for the probability to be PD1^+^ T cell versus PD1^−^ T cell show strong positive effects of cancer and macrophages and negative effects of T cells and fibroblasts.

For both cell types, we computed the probability that a cell is in each state as a function of the neighborhood composition of cell types. We used logistic regression, with all cell types in the system (B cells, T cells, Endothelial cells, Macrophages, cancer cells). The probability of each state varies across the different neighborhoods in the spatial tumor samples by a factor of 3 for T cells and 2.5 for fibroblasts (Fig 2a,b). We find that the cell state strongly associates the neighborhood composition regression model (p<<10^-10), as opposed to a control in which cell states were shuffled (p>0.1). We conclude that the cell type composition of the neighborhood is a strong correlate of the cell state.

Analyzing the regression coefficients allows us to estimate which cell types in the neighborhood contribute most strongly to the state probabilities. CAFs are favored in environments with T cells, endothelial cells and tumor cells (the latter are abundant-47% of the cells in the data- and have a large effect), Fig 2c). PD1 positive T cells are favored over PD1 negative T cells in environments with macrophages and tumor cells (Fig 2d).

### OSDR2.0 separates responders and nonresponders to chemotherapy and immunotherapy in triple negative breast cancer

We tested whether OSDR2.0 can discriminate between populations of responders and non responders to cancer therapy. We used the dataset from Wang et al (Wang *et al*., 2023) on the response of TNBC to two forms of treatment - chemotherapy and chemotherapy+immunotherapy (PD1 checkpoint inhibition). Each patient has three biopsies analyzed by IMC - before, three weeks after treatment, and 6 months after treatment (Fig 4A). This design provides a responder/non-responder status to each patient according to the post-treatment datapoint.

We trained OSDR2.0 on the responder population and the nonresponder population using the biopsy samples three weeks after treatment, and simulated the dynamics till cancer reaches steady state (∼100 days). One consideration is which initial condition to use in the simulations. We find that initial conditions do not affect the results - e.g. using initial conditions from responders, non-responders, or both yielded the same conclusions. Thus the dynamic rules (learned from responders or non-responders) rather than the initial condition seem to be dominant.

We studied OSDR2.0 with two types of cell state transition - transitions between PD 1 and PD1 T cells, or transitions between fibroblasts and CAFs defined by SMA, PDGFRB and Podoplanin (PDPN) as in (Danenberg *et al*., 2022) (Lavie *et al*., 2022; Cords *et al*., 2023). The data size (number of cells per patient) did not allow modelling both transitions simultaneously (Methods).

We simulated the models and tracked cancer cell numbers. We find that OSDR2.0 models trained on responders showed a reduction in cancer cells compared to the models trained on non-responders (Fig 4B). In response to immunotherapy (PD1 blockade+chemotherapy), OSDR2.0 shows higher peak levels of PD1+ T cells in responders than non-responders (Fig 4C).

To test the ability of ODSR2.0 to detect responders and non-responders from the biopsy taken during early treatment, we trained the model 100 times using subsampling of patients with returns. OSDR2.0 separates responders and non responders well, according to the criterion of rising or falling of cancer cell counts (Fig 4D). Separation is good in both chemotherapy and immunotherapy+chemotherapy arms (AUC>=0.96, Fig 3A).

**Figure 3.**
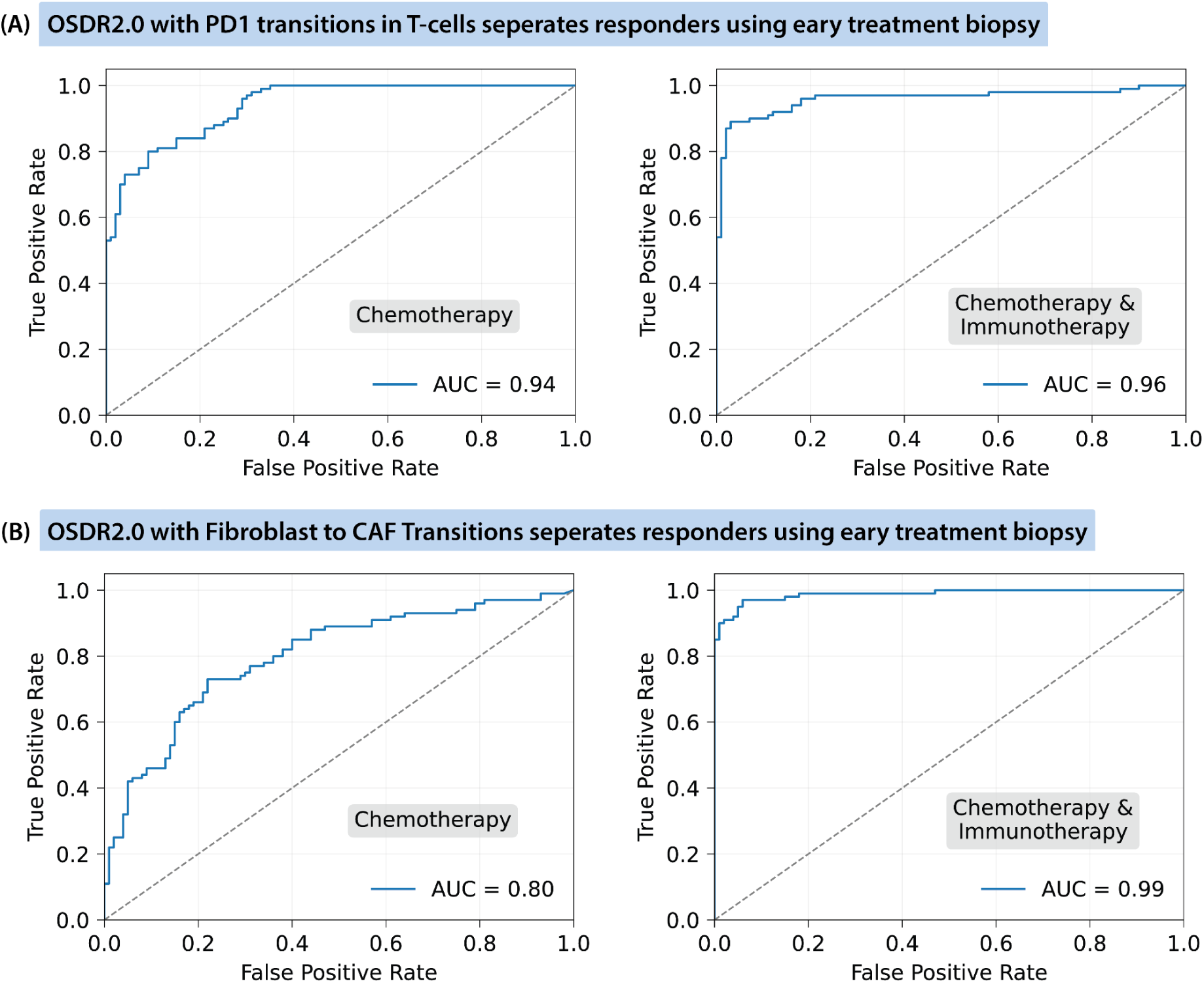
OSDR2.0 predicts responders to treatment from TNBC early-treatment biopsies. Receiver Operating Characteristic (ROC) curves (true-positive rate vs. false-positive rate) for discrimination between responders and non-responders; the diagonal is chance performance (AUC = 0.5) and AUC = 1.0 indicates perfect separation. (A) Model trained on PD1 transitions in T cells: chemotherapy only (left; AUC = 0.94) and chemotherapy + immunotherapy (right; AUC = 0.96). (B) Model trained on fibroblast to CAF (SMA, PDGFRB,PDPN) transitions: chemotherapy only (left; AUC = 0.80) and chemotherapy + immunotherapy (right; AUC = 0.99).

We asked what features enable OSDR2.0 to separate responders and non-responders in the rule set that it learns. The main difference picked up by OSDR2.0 is that cancer cells with abundant T cells in their neighborhood have a lower division probability in responders. In chemotherapy, this effect was due primarily to PD1 T cells, as evidenced by very low proliferation of cancer cells in the 10% environments most enriched with PD ^+^ T cells (Fig 4E).

**Figure 4.**
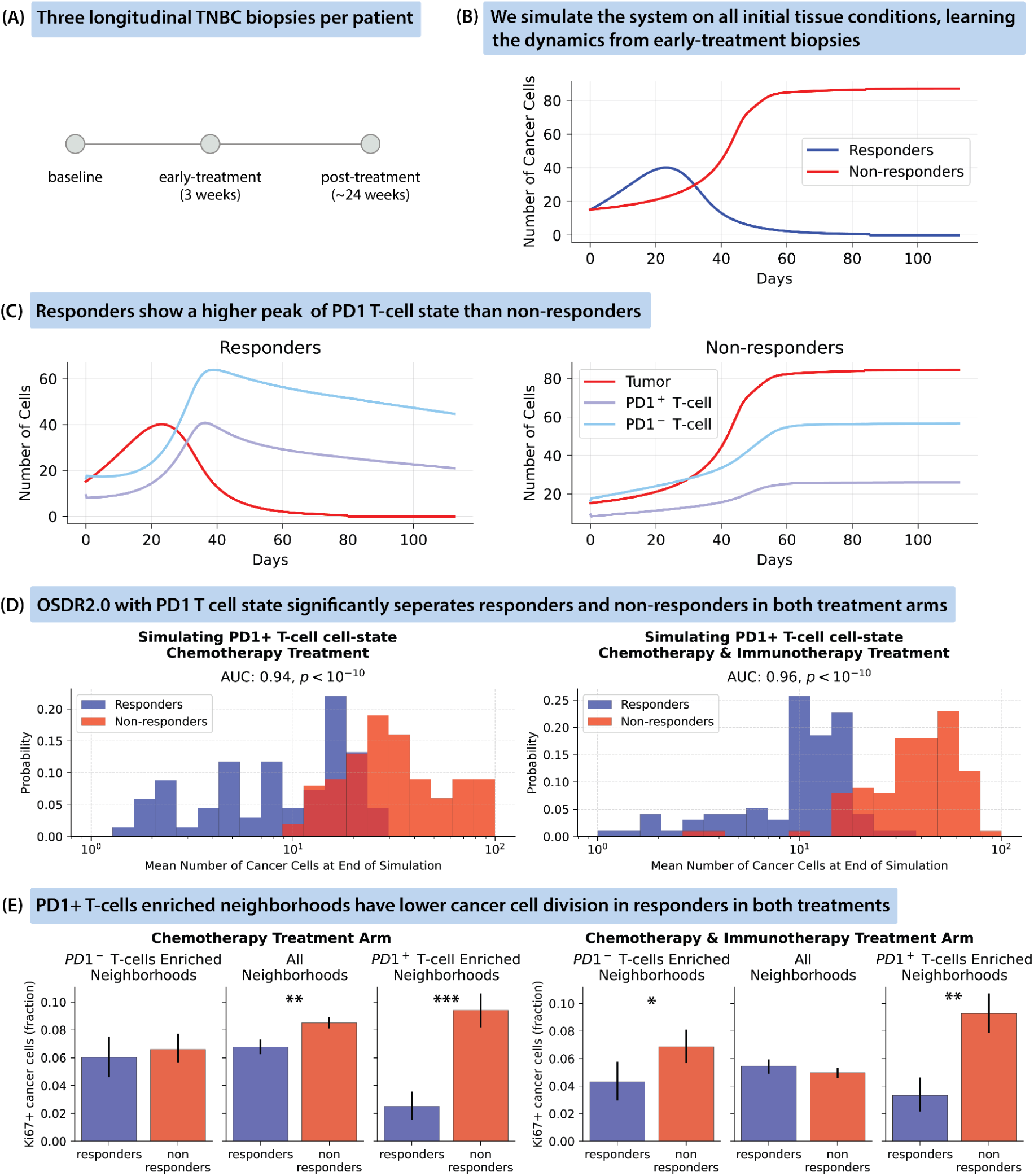
OSDR2.0 identifies responders to immunotherapy and chemotherapy in triple negative breast cancer. (A) The Wang et al dataset (Wang et al., 2023) has three longitudinal biopsies per triple negative breast cancer (TNBC) patient (𝑛 = 618). (B) OSDR2.0 with transitions between PD1+ T cells and PD1-T cells was applied to early-treatment 3w biopsies to learn cell-cell interaction dynamics for the responder and non responder participants (according to ∼24w biopsy). Simulations initialized with early-treatment conditions (𝑡 = 0) revealed divergent growth trajectories for responders and non-responders. (C) In responders OSDR2.0 shows higher peak levels of PD1+ T cells than in non-responders. (D) OSDR2.0 trained on responders shows fewer cancer cells at the end of the simulation than OSDR2.0 trained on non-responders under both chemotherapy (left) and combined chemo-immunotherapy (right) arms. (E) In PD ^+^ T-cell enriched neighborhoods, cancer cell division (Ki67 positivity) was lower in responders compared to non-responders across both treatment arms. Error bars are 95% confidence intervals. *𝑝 < 0. 05, ** ^−5^, *** ^−10^.

We also simulated OSDR2.0 with transitions between fibroblasts and CAFs (Fig 5). We again find that OSDR2.0 trained on responders shows a drop in cancer count over months, whereas training on non responders leads to rising cancer counts (Fig 5A). The CAF population shows a transient rise in CAFs that drops when the cancer count drops. In contrast non responders show a persistent CAF population (Fig 5A). The algorithm separates responders and non responders well (Fig 4D, 5B) especially in the chemotherapy+immunotherapy arm (Fig 3). We find that responders differ from non-responders in the proliferation of T cells - responders had higher T cell proliferation in CAF-rich neighborhoods than non responders (Fig 5C).

**Figure 5.**
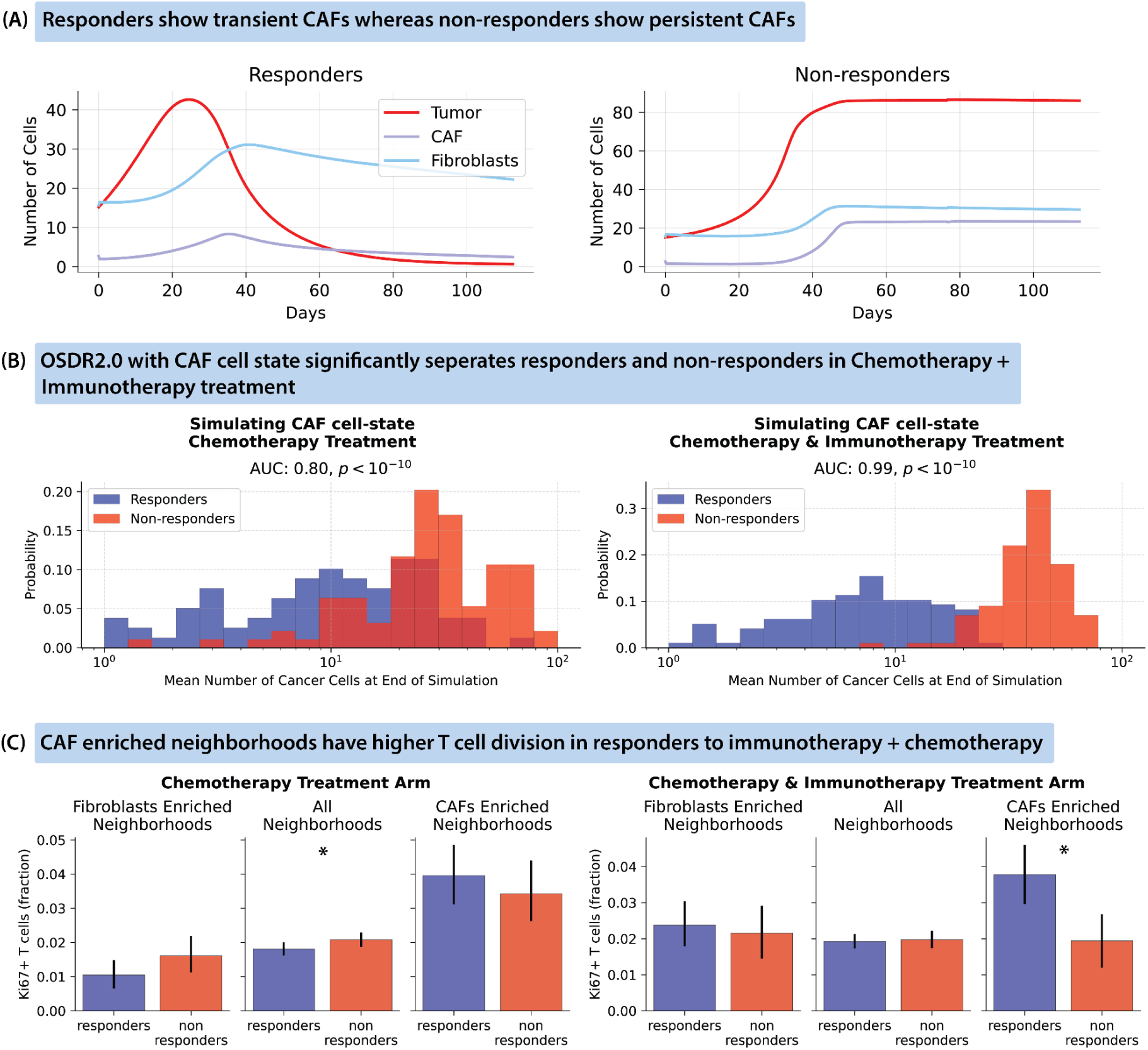
OSDR2.0 with fibroblast to CAF transitions identifies responders to immunotherapy and chemotherapy in triple negative breast cancer. (A) OSDR2.0 trained on responders showed a drop in cancer cells and a transient peak of CAFs, whereas OSDR2.0 trained on non-responders showed a rise in cancer and persistent CAFs. (B) OSDR2.0 trained on responders shows fewer cancer cells at the end of the simulation than OSDR2.0 trained on non responders. (C) In CAF-enriched neighborhoods, responders to Immunotherapy + Chemotherapy showed higher T-cell division under the combination treatment arm, suggesting a pro-immunogenic remodeling of the tumor microenvironment. Error bars are 95% confidence intervals. *𝑝 < 0. 05.

Simulations also include the immune cell types in the dataset (T cells, B cells and macrophages).

## Discussion

We developed OSDR2.0, a spatial modeling framework that extends OSDR1.0 by incorporating transitions between cell states within each cell type, based on local neighborhood composition. OSDR1.0 predicts cell population dynamics over weeks from a biopsy by learning division rates from neighborhood composition; OSDR2.0 adds to this prediction of cell states within a cell type. Using this approach, we demonstrate that cell states, such as CAF versus fibroblast and PD ^+^ versus PD ^−^ T cells are strongly associated with the composition of surrounding cell types in breast cancer biopsies. The model accurately captures these associations using logistic regression and simulates dynamic changes in both cell populations and cell states over time. Applied to data from triple-negative breast cancer (TNBC) patients undergoing chemotherapy or chemo-immunotherapy, OSDR2.0 simulations trained on responders show reduction in cancer cells over weeks, whereas OSDR2.0 trained on non responders show cancer persistances. These results suggest that a single tissue snapshot can provide dynamic information on cell populations and cell states over weeks.

OSDR2.0 generates a mathematical model for the effect of neighborhoods on cell division and state. This model can be used to detect which cell states in the neighborhood are most important for supporting cancer. In TNBC, the model suggests that the effect of PD1^+^ T cells on cancer cell division is key to response in both chemotherapy and immunotherapy. In both treatments, responders differed from nonresponders (at the 3 week treatment timepoint) by having slower cancer divisions in neighborhoods with high PD1^+^ T cells.

In order to employ OSDR2.0, one requires more cells per patient than for OSRD1.0, because OSRD2.0 has more parameters to describe cell states. The number of parameters rises quadratically with the number of cell types and states. Here we pooled responders and non responders to reach sufficient numbers of cells. We anticipate that emerging spatial technologies will provide a sufficient number of cells to allow analysis of individual patients. We estimate that on the order of 10,000 cells per cell type are needed depending on the probability of cell division and the frequency of each cell state.

One limitation is that OSDR2.0 does not account for migration of cells. Many cell types do not migrate appreciably over weeks (Fan and Rudensky, 2016; Houthuijzen *et al*., 2023; Suchanek *et al*., 2023). Others may be expected to migrate (such as immune cells) (Luster, Alon and von Andrian, 2005; Du *et al*., 2022). Since immune migration often involves vasculature (Xu *et al*., 2024), we expect that high migration would lead to positive coefficients for endothelial cells in the neighborhood of immune cells. Interestingly, we do not find strong positive coefficients for endothelial cells. We thus tentatively conclude that the TME we study here does not involve extensive migration of T cells, monocytes and other immune cells (developed TNBC). Future work can address migration, for example by utilizing markers for cell motility or chemotaxis.

In summary, OSDR2.0 offers a framework for inferring population dynamics and cell state transitions from a spatial snapshot of human tissue. By leveraging the strong association between cell state and neighborhood composition, the model provides a dynamic view of tissue organization and plasticity that was previously inaccessible from static data. Its ability to accurately separate responders from nonresponders in TNBC, based on simulations trained only on early post-treatment biopsies, highlights its potential as a predictive tool for treatment outcomes. Beyond oncology, OSDR2.0 can be applied broadly to spatial omics data across tissues and diseases, opening avenues for understanding microenvironmental control of cell states and guiding precision medicine.

## Methods

The number of cells of a given type in a tissue region can change due to cell division, cell death, migration (flux) or transdifferentiation into other cell types (Weinreb *et al*., 2018). In OSDR1.0 (Somer, Mannor and Alon, 2025), we focused on modeling cell-types which do not transdifferentiate at appreciable rates and analyzed tissues in which flux is negligible compared to rates of death or division (or alternatively, where flux terms can be added post-hoc). Under these assumptions, the population dynamics of cells of type 𝑖 are described by:

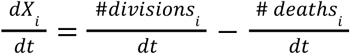

In OSDR2.0, we extend this model to include transitions between cell states within a cell type (e.g. fibroblast to CAF, PD1^−^ to PD1^+^ T cell). Letting 𝑋_*i*_ denote a specific cell type and 𝑋_*i*,α_ its subpopulation in state α, we update the equation to include transitions between these internal states:

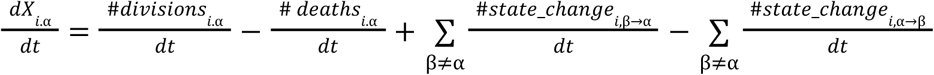

To handle the difference in timescales between slow processes (division and death; timescale T, on the order of weeks) and fast state transitions (timescale t, on the order of hours), we rewrite the above in terms of these separated scales:

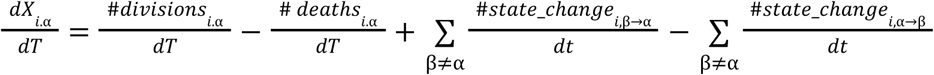

Because state transitions typically occur much faster than the cell division or death, we assume a quasi-steady state for the fast dynamics. From the perspective of the slow dynamic timescale T, this implies:

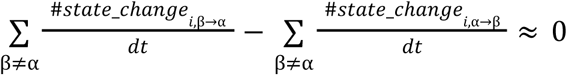

That is, at each time T, the distribution of cell states rapidly equilibrates to a steady-state that is determined by the local microenvironment.

A cell type’s rates of division, death, and state transitions are influenced by a range of local factors including, intercellular signaling, access to nutrients, genetics, and direct contact with neighboring cells. We collectively refer to this context as the cell’s neighborhood, and the set of features used to approximate it is denoted 𝑁(𝑥) for a cell 𝑥.

Under the quasi-steady state assumption, and given a spatial snapshot of the tissue, we can learn the probability that a cell is in a given state α, denoted 𝑝_α_ (𝑁(𝑥)) directly from its neighborhood:

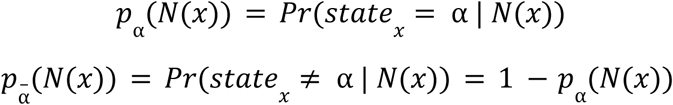

In OSDR2.0, this probability is modeled using logistic regression based on the composition of cell-types and states as 𝑁(𝑥), the set of features used to approximate the neighborhood of a cell, as detailed in the following sections.

OSDR1.0 estimates the expected change (𝑑𝑖𝑣𝑖𝑠𝑖𝑜𝑛 − 𝑑𝑒𝑎𝑡ℎ) rate of a cell based on its neighborhood, 𝔼[𝑥̇ | 𝑁(𝑥)], without accounting for cell state heterogeneity. In contrast, OSDR2.0 models the probability of a cell being in a specific state α, 𝑝_α_ (𝑁(𝑥)), which allows to decompose the expected rate of change (𝑥̇) as:

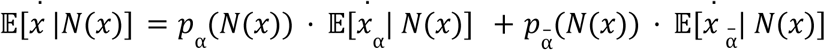

More generally, if 𝑆_*t*[*x*]_ denotes the set of all possible cell states for the cell type 𝑡[𝑥] of a cell 𝑥, we can write:

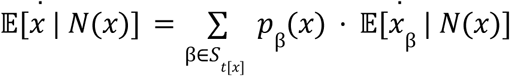

This formulation shows that OSDR2.0 extends OSDR1.0 by explicitly modeling cell state distributions, enabling more accurate estimation of functional behaviors of cell states.

To implement OSDR2.0, we first stratify each cell type into its defined states (e.g., PD1^+^ and PD1^−^ T cells), and then apply OSDR1.0 independently to each state as if it were a separate cell type. This yields division and removal dynamics for each state-conditioned subpopulation. We then rebalance these dynamics at each simulation step according to the OSDR2.0-estimated state probabilities from the current neighborhood, ensuring that state transitions remain in quasi-equilibrium. This separation enables OSDR2.0 to simulate realistic state-dependent population dynamics from static tissue data.

Like OSDR1.0, OSDR2.0 is agnostic to the choice of inference model: while we use logistic regression in this work, other statistical or machine learning models can be integrated within the framework. Moreover, the approach is compatible with any spatial proteomics technology that enables accurate classification and localization of individual cells, along with robust measurement of markers for division and cell states.

## Model Inference Algorithm

The following algorithm estimates 𝑝_α_, the probability that a cell is in a particular state α

Input:

A set of tissues represented as the ordered set (𝐵, 𝑇, 𝑙, 𝑆, 𝑂), a set of cell IDs {1, ···, 𝑁} and a neighborhood radius 𝑟 :

● 𝐵 - a set of tissues, each containing a subset of cell IDs.
● 𝑇 - a mapping from a cell 𝑥 to its cell type 𝑇[𝑥], for simplicity we also denote by 𝑇 the set of all cell types..
● 𝑙 - a mapping from a cell 𝑥 to its spatial coordinates in the tissue 𝑙[𝑥].
● 𝑆 - a mapping from each cell type 𝑡’ ∈ 𝑇 to its possible set of cell states, for + − {𝑃𝐷1^+^ 𝑇𝑐𝑒𝑙𝑙, 𝑃𝐷1^−^ 𝑇𝑐𝑒𝑙𝑙} 𝑡’ = 𝑇𝑐𝑒𝑙𝑙. In this algorithm we assume |𝑆[𝑡’]| = 2 𝑜𝑟 |𝑆[𝑡’]| = 1 for simplicity, where if |𝑆[𝑡’]| = 2 then 𝑆[𝑡’] = {α, ᾱ } and if |𝑆[𝑡’]| = 1 then the cell has only one cell state.
● 𝑂 - a mapping from a cell 𝑥 to its observed cell-state 𝑂[𝑥] ∈ {0, 1} indicating whether 𝑥 is in cell state α or not.

Algorithm:

1. For each tissue 𝐵_*i*_ ∈ 𝐵 do 1.1. For each 𝑥 ∈ 𝐵_*i*_ do 1.1.1. If α ∉ 𝑆[𝑇[𝑥]] skip 𝑥 1.1.2. For each cell type 𝑡’ ∈ 𝑇 do 1.1.2.1. Compute the number of neighboring cells of 𝑥 of type t’

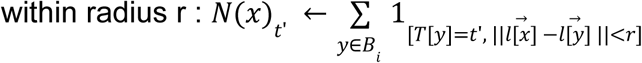
2. Apply feature transformation to 𝑁(𝑥) such as adding polynomial features and interaction terms, yielding the transformed feature vector 𝑋(𝑥).
3. Use the transformed features 𝑋(𝑥) and 𝑌(𝑥) = 𝑂(𝑥) as labels to fit a multivariate logistic regression model (or any other suitable model), denote it *p*_α_

Output:

Return the fitted model 𝑝_α_.

## Model Simulation Procedure

Let

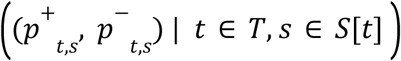

denote the division and death probabilities for all cell types 𝑡 in all states 𝑠, obtained by applying OSDR1.0 model inference while treating each cell state as a distinct cell type.

In OSDR1.0, simulation can be ran by iteratively applying division and death probabilities for each cell 𝑥 according to its type 𝑇[𝑥] and neighborhood 𝑁(𝑥), i.e., using 𝑝^+^_*T*[*x*]_ (𝑁(𝑥)), 𝑝^−^_*T*[*x*]_ (𝑁(𝑥)), In OSDR2.0 we also have cell state so we will use 𝑝^+^_*T*[*x*]*S*[*x*]_ (𝑁(𝑥)), 𝑝^−^_*T*[*x*],*S*[*x*]_ (𝑁(𝑥)) instead.

In OSDR2.0, an additional rebalancing step is required: after simulating division and removal, the distribution of cell states must be adjusted according to the predicted state probabilities 𝑝_α_ from the OSDR2.0 model inference process.

To make this process interpretable and consistent, we simulate dynamics on a phase portrait where the axes represent the number of cells in each state-specific subtype (Figure 1). After each application of the OSDR1.0-based dynamics, the population is rebalanced so that, for each cell type, the proportion of cells in each state aligns with the predicted probabilities using the model 𝑝_α_. This ensures that the cell state distribution remains consistent with the inferred steady-state dynamics.

## Data Analysis

We used the datasets from Wang et al (Wang *et al*., 2023) and Danenberg et al (Danenberg *et al*., 2022), we used their cellular typing, definitions, reducing them to more general cell types, and added cell-states per analysis.

The cell types used were: Fibroblasts, T-cells, B-cells, Macrophages, Cancer cells and Endothelial cells.

The phenotypes used in the analyses were PD ^+^ versus PD ^−^ T-cells and CAFs versus non CAF Fibroblasts.

